# Shared and diverging neural dynamics underlying false and veridical perception

**DOI:** 10.1101/2023.11.16.567367

**Authors:** Joost Haarsma, Dorottya Hetenyi, Peter Kok

## Abstract

We often mistake visual noise for meaningful images, which sometimes appear to be as convincing as veridical percepts. This suggests that there is considerable overlap between the mechanisms that underlie false and veridical perception. Yet, false percepts must arise at least in part from internally generated signals. Here, we apply multivariate analyses to human MEG data to study the overlap between veridical and false perception across two discrete stages of perceptual inference: discrimination of content (what did I see) and detection (did I see something?). To this end, participants performed a visual discrimination task requiring them to indicate the orientation of a noisy grating, as well as their confidence in having seen a grating. Importantly, on 50% of trials no gratings were presented (noise-only trials). On a subset of these noise-only trials, participants reported seeing a grating with high confidence, dubbed here false percepts. We found that a sensory signal reflecting the content of these false percepts was present both before and after stimulus onset. Uniquely, high confidence false, but not veridical, percepts were associated with increased pre-stimulus high alpha/low beta [11-14Hz] power, potentially reflecting enhanced reliance on top-down signalling on false percept trials. Later on, a shared neural code reflecting confidence in stimulus presence emerged for both false and veridical percepts, as revealed by cross-decoding. These findings suggest that false percepts arise through sensory-like signals reflecting both content and detection signals, similar to veridical percepts, with an increase in pre-stimulus alpha/beta power uniquely contributing to false percepts.

**Significance statement:** The neural mechanisms underlying false percepts are likely different from those that underlie veridical perception, as the former are generated endogenously, whereas the latter are the result of an external stimulus. Yet, false percepts often get confused for veridical perception, suggesting a converging mechanism. This study explores the extent to which the mechanisms diverge and converge. We found that both high confidence false and veridical percepts were accompanied by content-specific stimulus-like orientation signals, as well as a shared signal reflecting perceptual confidence. In contrast, we found that false, but not veridical, percepts were preceded by increased high alpha/low beta [11-14 Hz] power, possibly reflecting a reliance on endogenous signals.

## Introduction

When we try to make sense of our noisy visual surroundings, we sometimes mistake our own internally generated signals for externally caused sensations. For example, a cat owner may have such a strong expectaion to see their pet that they misperceive a blurry flash in their visual periphery for it. Despite these false perceptual experiences relying on internally generated signals, they sometimes appear to be just as real as veridical percepts, suggesting at least partially overlapping neural mechanisms between veridical and false perception. However, the extent to which the underlying neural mechanisms are similar remains unclear.

How does the brain make inferences about the world? Perceptual inference has two main aspects: what is the content of the experience, and how sure am I that I perceived it? When a silhouette appears in the dark, we need to infer what it is (a tree, a person?) as well as make a decision on how likely it is that I perceived this object. Indeed, some recent empirical and theoretical work has highlighted the importance of separating these distinct and discrete stages of inference (Mazor et al., 2020a,b; Fleming et al., 2020; Dijkstra et al., 2023). The neural signals underlying false percepts may resemble those of veridical perception in both these aspects, i.e., there are signals reflecting the content of false percepts as well as signals reflecting confidence in having perceived said object. Alternatively, false percepts may not rely on sensory content signals, and only share signals reflecting confidence in having perceived something, rather than nothing. This important distinction also pertains to one of the most influential models of hallucinations, namely the predictive coding account of hallucinations. The predictive coding model states that hallucinations arise through increased influence of prior expectations (Powers et al., 2017; Kafadar et al., 2020; Haarsma et al., 2020; Benrimoh et al., 2023; Schmack et al., 2021). However, it remains unclear whether these priors act on the content level of stimulus discrimination, on the level of stimulus detection, or both. Therefore, characterising at which stages of perceptual inference false percepts arise is of crucial importance to further our understanding of hallucinations, as well as subjective perception more generally.

While false and veridical percepts may share considerable overlap, their neural origins must be separate in some sense, as they rely on different neural inputs by definition – the former being generated internally, and the latter externally. Pre-stimulus oscillatory dynamics are a candidate mechanism for a point of divergence. Previous studies have consistently linked a decrease in alpha power to increased hit rates by inducing a more liberal detection threshold, while not affecting sensitivity, possibly through increasing neuronal excitability (Achim et al., 2013; Benwell et al., 2017; Boncompte et al., 2016; Brüers & VanRullen, 2018; Chaumon & Busch, 2014; Ergenoglu et al., 2004; Iemi et al., 2017; Iemi & Busch, 2018; Mathewson et al., 2014; Samaha et al., 2017, 2020; Van Dijk et al., 2008). Yet, evidence linking decreased alpha to increased false percepts remains mixed (Keil et al., 2014; Lange et al., 2013). In contrast, a second body of work has linked increased alpha and beta to top-down cognitive processes believed to contribute to false percepts. For example, increased beta power plays a role in interpreting ambiguous stimuli in the language (Iversen et al., 2009) and visual domains (Hipp et al., 2011; Okazaki et al., 2008), as well as mental imagery (Bartsch et al., 2015; Villena-González et al., 2018), and perceptual predictions (Auksztulewicz et al., 2017; Bastos et al., 2012, 2020; Engel et al., 2001; Fujioka et al., 2012; Sherman et al., 2016; Turner et al., 2023). Thus, if top-down mechanisms play a role in driving false percepts, we might find enhanced power in the alpha/beta frequency bands to be related to false percepts.

Here, we used multivariate decoding techniques in combination with MEG to study the overlap between false and veridical perception at the content and detection-level, as well as to study a potentially unique role for neural oscillations in driving false percepts. To preview, false percepts resembled veridical perception to the extent that both were associated with stimulus-like content-specific signals reflecting the perceived gratings, as well as a shared signal reflecting confidence in stimulus presence. Uniquely however, false, but not veridical percepts were associated with an increase in pre-stimulus high alpha/low beta (11-14Hz) power, possibly reflecting endogenous feedback signals. Together, these findings show that false percepts arise from sensory-like neural mechanisms, similar to veridical perception.

## Results

### Participants experienced false percepts that were independent of perceptual expectation cues

Participants performed a grating discrimination task under noisy conditions, while on 50% of the trials noise-only patches were presented. Participants indicated both the perceived orientation as well as their degree of confidence in having perceived a grating (regardless of orientation). Participants accurately identified the grating orientation on grating-present trials more often than expected by chance (mean accuracy=0.66, SD=.12, T{24}=6.5, *p*<.001). Furthermore, they were more accurate when they were confident that they had seen a grating (i.e., higher than average confidence across trials) than when they were not (high: mean=.70, SD=.13; low: mean=.59, SD=.11; paired t-test) (T{24}=6.76, *p*<.001) (Fig. 1c), demonstrating that they were able to perform the task and used the confidence ratings in a meaningful way. It is worth repeating that here participants reported their confidence in having seen a grating, rather than confidence in their orientation report, and thus effectively gave a perceptual awareness response (Sandberg & Overgaard, 2015). Participants were slightly more confident on grating-present trials (mean confidence =2.29, SD=.64, on a scale of 1-4) than noise-only trials (mean=2.25, SD=.64) (T{24}=2.28, *p*=.031). Upon debriefing, all participants but one underestimated the frequency of noise-only trials, believing on average that .18 (SD=.16) of trials contained just noise, while the true proportion was .50 (Fig. 1d). Strikingly, participants reported perceiving a grating with high confidence (3 out of 4 or higher) on 35% of noise-only trials (Fig. 1f). The perceptual expectation cues significantly biased which orientation participants perceived on noise-only trials (.55 false percepts congruent with the cue, chance level is .50, T{24}=2.53, *p* =.018). This effect was driven by the individuals who became aware of the meaning of the cues (N = 7 out of 25; Fig. 1e), potentially reflecting a response bias. Indeed, those aware of the cue had significantly stronger effects of the cue (T{24}=4.14, *p*<.001). High confidence false percepts were not more affected by the cues than low confidence percepts, i.e. guesses (T{23}= 1.18, *p*=.24). In sum, these results indicate that participants regularly had false grating percepts, but that these were hardly, if at all, affected by the predictive cues. This is highly consistent with a previous study using the same experimental paradigm (Haarsma et al., 2023).

**Fig. 1.**
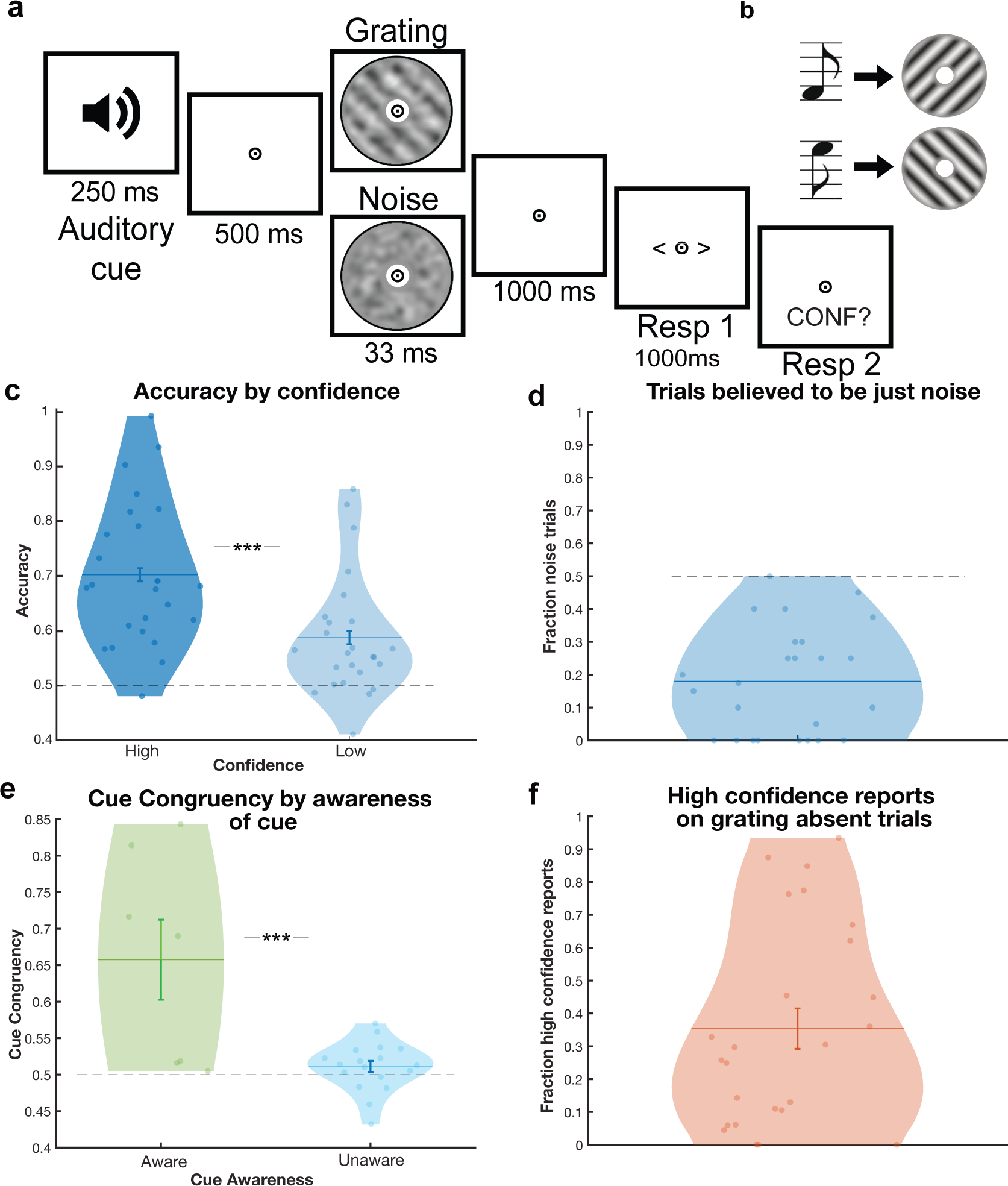
Experimental design and behavioural findings. **a**, An auditory cue was followed by either a low contrast grating embedded in noise (50% of trials), or a noise patch (50%). Participants indicated which orientation they saw and how confident they were that a grating was presented. **b**, One sound predicted the appearance of a 45°, or clockwise, oriented grating, whilst the other predicted a 135°, or anti-clockwise, orientated grating. Auditory cues were 100% valid on grating-present trials. **c,** Participants were more accurate on high confidence trials. **d,** Upon debriefing participants believed that only 18% of trials contained noise, compared to the real number of 50%. **e,** Participants were driven by the cue on noise-only trials when they were aware of the cues meaning. **f,** On noise-only trials, participants reported high confidence in having seen a grating on 35% of trials.

### Orientation-specific activity reflecting the perceived stimulus was present both pre- and post-stimulus on high confidence false percept trials

In order to test the hypothesis that false percepts arise from sensory-like processes we performed a linear discriminant analysis (LDA) to decode the orientation of high contrast, task-irrelevant gratings presented in a separate localiser block in order to then test for the presence of these signals during the main experiment. First, we confirmed that the orientation of these high contrast gratings could be decoded from the MEG signal (Fig. 2a). Cluster-based statistics on the diagonal of the temporal generalisation matrix revealed significant decoding from 55ms-270ms post-stimulus (*p*<.001, Cohen’s d= 1.87), peaking at 105ms (Fig. 2b). Further, source localisation revealed that the occipital lobe contributed primarily to the decoding of the grating orientation (Fig. 2c). We then generalised from the localiser to the main experiment to see whether sensory signals contributed to high confidence false percepts. Here, we trained the decoder on a 10ms window centred on the peak of the localiser decoding (100-110ms). We subtracted orientation-specific stimulus traces on low confidence trials from high confidence ones, for both grating present and noise-only trials. We focussed on the pre- and post-stimulus time-windows separately (−500 to 0ms and 0 to 1000ms, respectively). First, across all trials we found a clear orientation-specific neural signal for high vs. low confidence trials during the post-stimulus time-window, from 180ms-855ms (*p*<.001, Cohen’s d= 0.87) (Fig. 2d). Analysing grating-present and noise-only trials separately revealed orientation-specific signals for grating-present trials from 505ms-610ms and 865ms-940ms (*p*=.018 & *p*=.043, Cohen’s d= 0.75 & 0.71) (Fig. 2f), and critically also on noise-only trials from 390ms-510ms (*p*=.014, Cohen’s d= 0.68) (Fig. 2e). Further, during the pre-stimulus time window there was significant orientation-specific activity on noise-only trials from −55ms to −5ms & −205ms to −150ms (*p*=.047 & *p*=.050, Cohen’s d= 0.64 & 0.52) (Supplementary Fig.1). Note, because only noise patches with low (2%) signal energy for all orientations were selected to be included in the present experiment, this orientation-specific activity could not have arisen from the noise-patch itself. No pre-stimulus orientation-specific signals were found on grating present trials. Taken together, false percepts were associated with orientation-specific sensory-like signals in both the pre-stimulus and post-stimulus time windows.

**Fig. 2.**
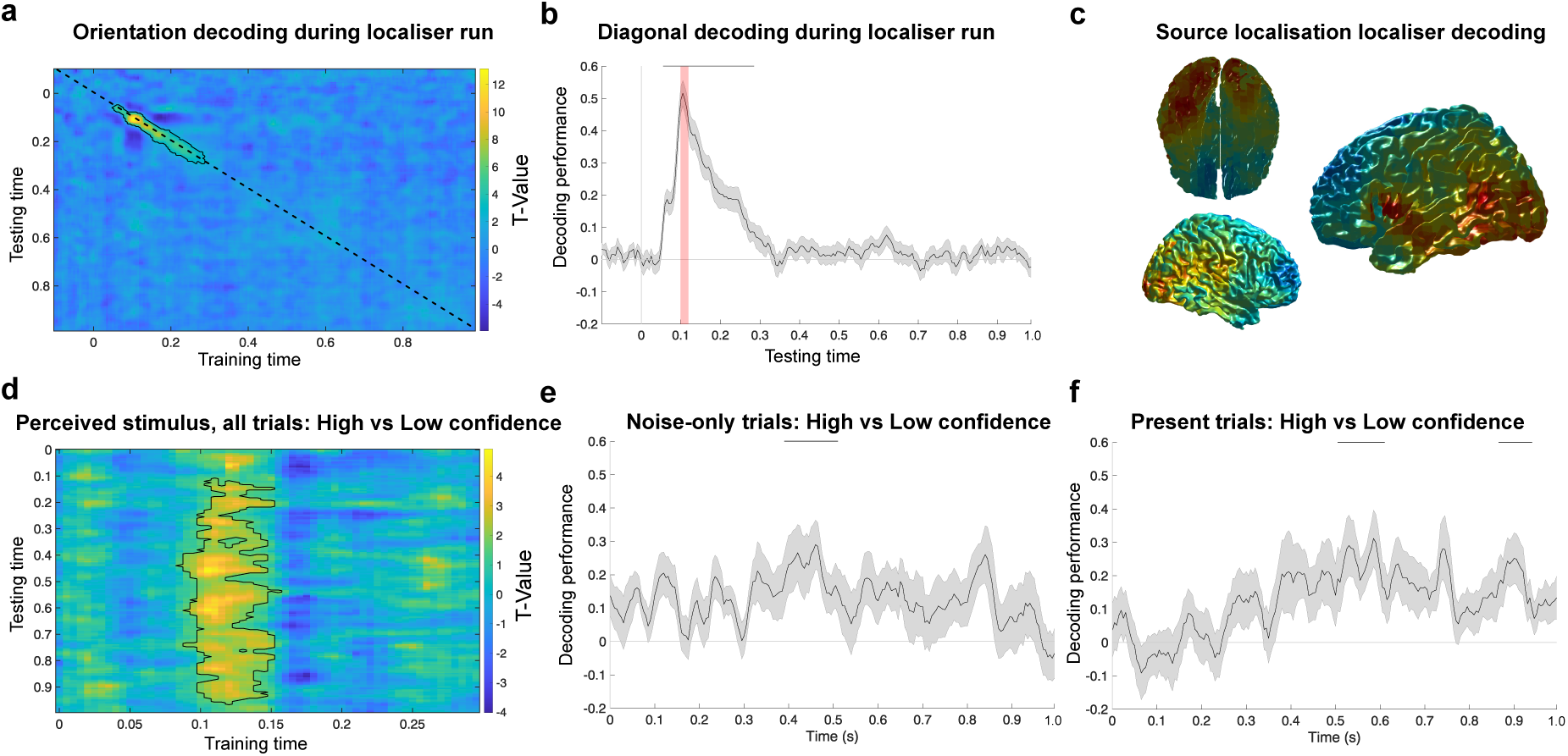
Decoding of perceived orientation. A linear discriminator analyses was used to decode orientation-specific activity from a functional localiser, which was then generalised to the main experiment. The orientation of a clearly presented grating could be decoded from the functional localiser with strong clusters in the temporal generalisation matrix **(a)** and the diagonal **(b). b,** The peak of the localiser window that served as the training timepoint for the generalisation to the main experiment is indicated in shaded red. **c,** Orientation decoding was primarily driven the occipital lobe. **d,** Generalising from the localiser to the main experiment revealed a clear signal reflecting the perceived orientation across all trials. Training on the peak of the localiser decoding (100-110ms) and separating by stimulus presence revealed significant clusters on noise-only trials **(e)** and stimulus present trials **(f).**

*A shared neural code reflecting confidence in stimulus presence emerged 250ms post-stimulus* We next decoded confidence in stimulus presence on both grating-present and noise-only trials using an LDA (Mostert et al., 2015). Rather than training on a separate localiser (during which participants did not give confidence responses), here we used cross-validation to train and test within the main experiment blocks, in order to isolate a signal reflecting participants’ confidence in having seen a grating (regardless of its content). Across both grating-present and noise-only trials, a confidence signal emerged 265ms post stimulus and was sustained throughout the post-stimulus window (265ms-2000ms, *p*<.001, Cohen’s d= 1.6) (Fig. 3a). This signal was present on both noise-only (410ms-1280, 1280-2000ms, *p*<.001, Cohen’s d= 1.1, 1.2) (Fig. 3b) and grating-present trials (425ms-670, 700-2000ms, *p*<.001, Cohen’s d= 0.83, 1.26) (Fig. 3c). To test whether the neural representation of perceptual confidence in veridical and false percepts was the same, we trained on perceptual confidence on noise-only trials, and decoded the confidence on grating-present trials, and vice versa. Cluster-based analyses revealed a significant effect in the post-stimulus time window, demonstrating that the confidence signal on noise-only trials generalised to grating-present trials (235ms-385ms, 415ms-470ms, 480ms-2000ms, *p*<.001, Cohen’s d= 0.85, 0.76, 1.38) (Fig. 3d) and vice versa (225ms-295ms, 320ms-385ms, 415ms-875ms, 885ms-1000ms, *p*=.008, .015, .001, .003, Cohen’s d= 0.93, 0.81, 1.26, 1.19) (Fig. 3e).

**Fig. 3.**
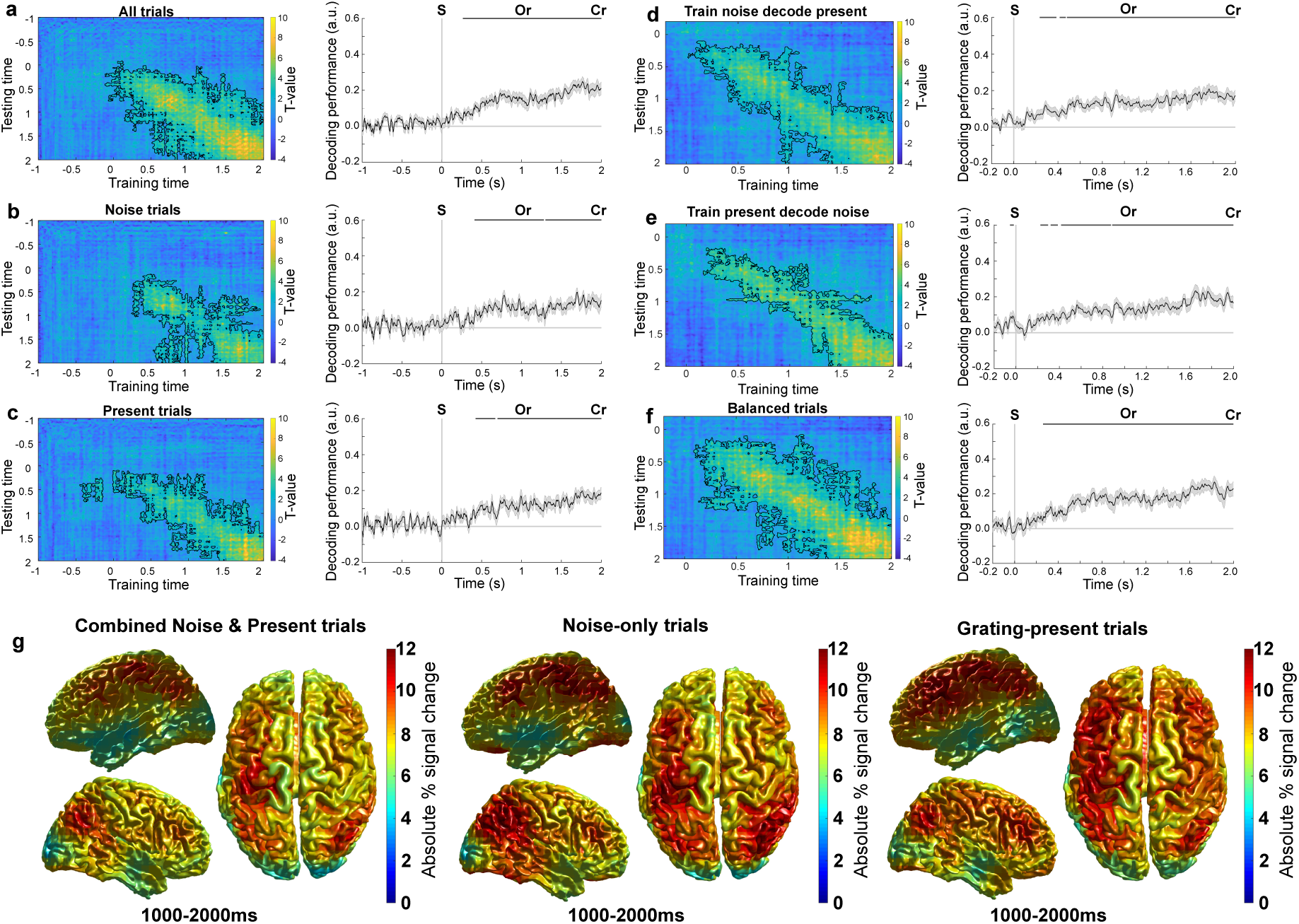
Decoding of perceptual confidence. A linear discriminator analyses was used to identify a perceptual confidence signal. **a,** A significant confidence signal was found on all trials after stimulus onset. Analysing noise-only **(b)** and grating-present **(c)** trials separately revealed similar results. Cross-decoding from noise-only to grating-present trials **(d),** and vice versa **(e),** demonstrated that there was a shared neural signal on grating-present and noise-only trials. **f,** When balancing low and high confidence trial counts decoding was still successful, demonstrating that the effect was not confounded by imbalanced trial counts. **g,** Percentage absolute signal change of the high confidence condition compared to the low confidence condition in source localisation. Here the combined, noise-only, and grating-present trials in the 1000ms to 2000ms time window are presented separately. Across all three conditions, parietal and frontal cortices contributed most strongly to the linear discrimination analyses. S = onset of stimulus, Or = onset orientation response cue, Cr = onset of confidence response cue.

To ensure that the decoding results were not confounded by unequal numbers of low and high confidence trials, we repeated the analyses drawing a random number of trials from the overrepresented condition equal to the number of trials in the underrepresented condition. In some individuals this led to very few trials to perform the decoding analyses on, leading to unreliable results. We therefore removed participants with fewer than 50 trials to train the decoder on. This control analysis confirmed significant decoding of confidence post-stimulus (all trials: 260ms-2000ms ; Fig. 3f).

For visualisation purposes, we reconstructed the cortical sources of the confidence signal by performing source localisation separately on the magnetic fields underlying high and low confidence perceptual reports and computing a signal difference map. This source reconstruction was conducted on the 1000-2000ms time window, where decoding was the strongest. The source map was overlaid on a 3D cortical surface, revealing parietal and frontal regions for both noise-only and grating-present trials (Fig. 3g).

### Within experiment cross-validated decoding of perceived orientation

Note that the confidence decoding analysis involved cross-validated training and testing within the main experiment, whereas for the orientation decoding analysis we trained on a separate localiser and generalised to the experiment in order to provide a strong test of the sensory nature of the orientation signals. For completeness’ sake, and to aid comparison with confidence decoding, we also applied a within experiment cross-validation approach to decoding the reported orientations (Supplementary Fig. 2). Broadly, this confirmed our finding that subjective percepts were reflected by orientation-specific neural signals that were modulated by confidence (Supplementary Fig. 2d). Note, these signals cannot reflect motor responses, as the response mapping was randomised. When decoding participants motor responses, a significant cluster was present exclusively in the response time-window (cluster-based analyses on diagonal, all trials: 1015-2000ms, *p*<.0001, Cohen’s D=2.43), which was not modulated by confidence (*p*>.1).

### False percepts were preceded by an increase in pre-stimulus high alpha/low beta power

To explore whether pre-stimulus neural oscillations played a role in driving false percepts, we conducted a frequency analyses on the pre-stimulus time window. Specifically, we tested whether changes in power in the alpha and beta bands preceded high confidence false percepts, that is, trials on which participants indicated high confidence in having seen a grating in the absence of one. We tested for an interaction using a two-way repeated measures ANOVA with confidence and stimulus presence as two-level factors within the - 750ms to 0ms time window (i.e., the time window from the auditory cue to the onset of the stimulus). There was a significant interaction in the beta band (12-20Hz) (−650ms to −50ms, *p*=.0039), but not in the alpha band (8-12Hz, −650ms to 0ms, *p*=.0939). Post-hoc analyses revealed that this interaction was driven by an increase in beta power preceding high confidence false percepts (−650ms to 0ms, *p*=.0108, cluster-based permutation-test) (Fig. 4a). We conducted post-hoc analyses to estimate the exact frequencies that drove these effects by repeating the analyses in steps of 1Hz. This revealed the interaction between stimulus presence and confidence started in the high alpha band (11Hz, −700ms to −300ms, *p*=.0499; 12Hz, −700ms to 0ms, *p*=.002) and continued into the beta band (13Hz, −700ms to 0ms, *p*=.0080; 14Hz, −650ms to −50ms, *p*=.030), as did the increase in beta power preceding high confidence false percepts (11Hz, −700ms to 0ms, *p*=.0039; 12Hz, −700ms to 0ms, *p*=.0059; 13Hz, −750ms to −150ms, *p*=.0079; 14Hz, −700ms to −400ms, *p*=.049).

**Fig. 4.**
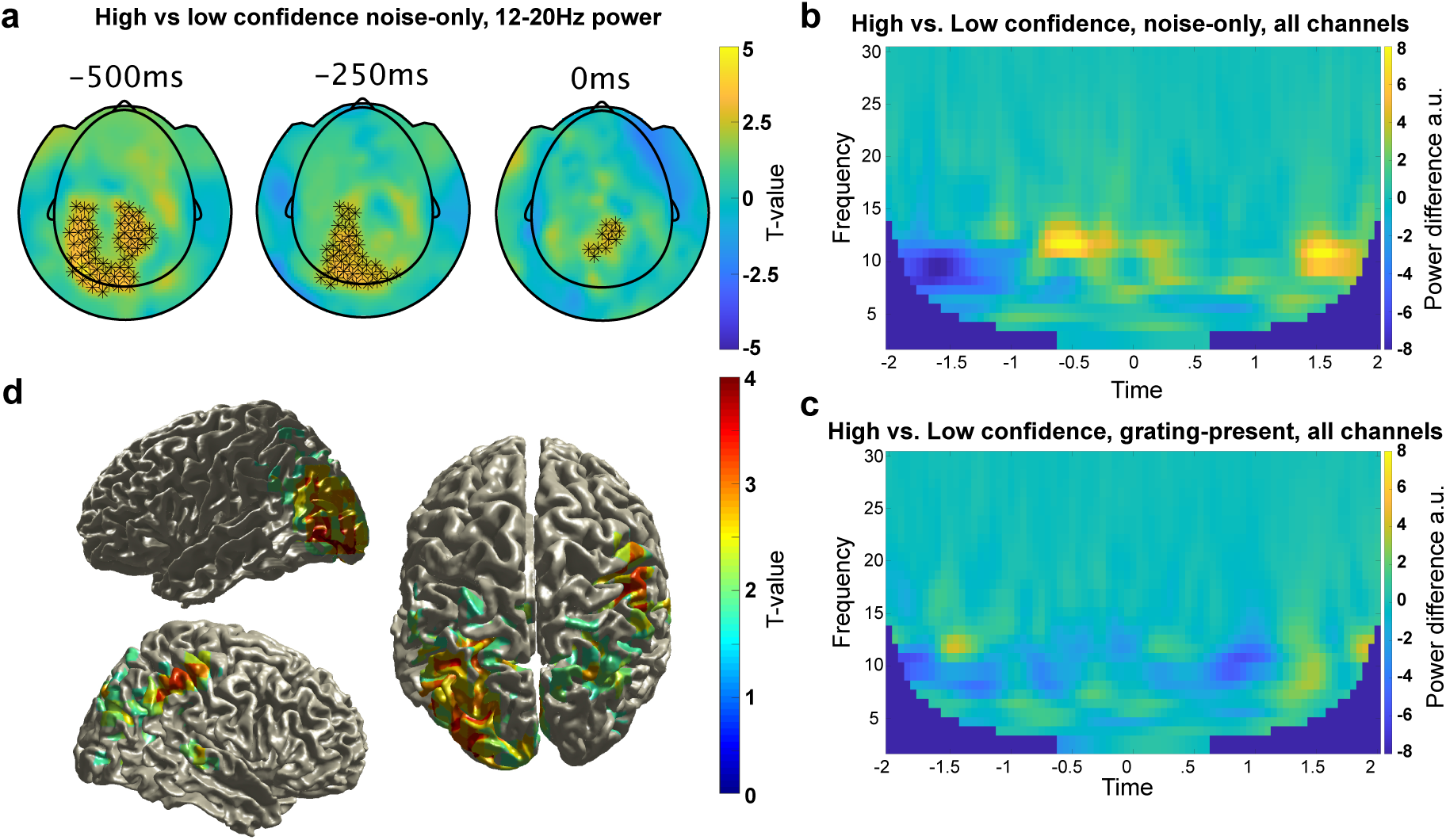
Frequency analyses. **a,** Scalp topographies of differences in pre-stimulus beta power for high minus low confidence false percepts. **b,** Difference in average time x frequency between high and low confidence trials on noise-only trials.. **c,** Difference in average time x frequency between high and low confidence trials on grating-present trials. **d,** T-map of the difference in source localisation for the 13Hz frequency band +- 1.5Hz between high and low confidence response on noise-only trials. A network of parietal and occipital regions was active prior to high confidence false percepts

Thus, the increase in power preceding high confidence false percepts was in the high alpha, low beta band (Fig. 4b). In contrast, no effects were found comparing high and low confidence reports on grating-present trials for either alpha or beta power, *p*>.5, Fig. 4c). No effects were found in the post-stimulus time-window for either false or veridical percepts (*p*>.5). In sum, an increase in high alpha/low beta band power preceded high confidence false percepts, but not high confidence veridical percepts.

To further explore the temporal profile of this effect, we extended the time window to - 2000ms to 0ms. This control analysis revealed that the interaction between stimulus presence and confidence on beta power started after the onset of the auditory cue (−650ms to −50ms, *p*=.018).

We performed source localisation analyses on the difference in beta power prior to high and low confidence false percepts in the −750 to 0ms pre-stimulus time window centred on 13Hz (+- 1.5 Hz). This revealed that the increase in beta power arose from a network including parietal and occipital regions (Fig. 4d).

### ERF analyses

Finally, we performed ERF analyses to explore whether confidence in stimulus presence was reflected in the amplitude of the ERFs on grating-present and noise-only trials. For these purposes we contrasted high and low confidence trials, and computed cluster-based analyses on these differences. There were no effects of confidence on ERF amplitude on the sensor level on either noise-only or grating-present trials (*p*>.5).

## Discussion

The present study investigated whether the neural mechanisms underlying false and veridical perception are shared. Specifically, we tested at which stage of perceptual inference commonalities between veridical and false perception emerge. During a perceptual discrimination task, participants indicated whether they perceived a grating with a specific orientation. On a subset of noise-only trials, participants reported seeing gratings with high confidence, which we dubbed high confidence false percepts. We found that on these false percept trials a sensory-like signal reflecting the perceived orientation emerged both prior to and following the onset of the noise stimulus. The same was true for veridical trials, albeit only post-stimulus. Further, a signal reflecting confidence in having seen a grating emerged around 250ms on both false percept and veridical trials, originating from a frontal/parietal network. Cross-decoding between false and veridical trials showed that the neural patterns underlying this confidence signal was shared. Finally, uniquely to high confidence false percept trials, there was an increase in pre-stimulus high alpha/low beta power (11-14Hz), arising from a parietal-occipital network. Together, these findings show that the neural signals that underlie false and veridical percepts are largely shared both on the level of sensory-like content signals and in terms of neural signals reflecting confidence in stimulus presence, with an additional unique role for pre-stimulus oscillations on false percept trials.

The present study distinguished two aspects of perceptual inference: content discrimination and detection. Recent theoretical work has highlighted the importance of these different stages of inference, and how they are conceptually different (Mazor et al., 2020; Fleming et al., 2020; Dijkstra et al., 2023). When applied to false percepts, this conceptual distinction suggests that these experiences could in principle arise in different ways. That, is a false percept could occur due to sensory-like activity (a sensory signal resembling a particular stimulus causes the agent to report a false percept), or through the neural mechanisms that are concerned with reporting the presence of a stimulus (I saw something rather than nothing). Here we find evidence suggesting that false percepts arise from low-level sensory-like signals reflecting the content of these false percepts. To our knowledge, this is the first study to decode the content of false percepts from the MEG signal and show that they resemble sensory signals. Two fMRI studies have reported content-specific signals related to false percepts. First, Pajani et al. (2015) linked spontaneous pre-stimulus orientation-specific activity to falsely reported orientations in noise. Second, a previous layer-specific fMRI study from our group used the same experimental paradigm as the present study (Haarsma et al., 2023). Notably, the behavioural findings across these two studies were highly consistent, with both reporting a limited effect of cued orientations on perception, which was only significant in those who became aware of the meaning of the auditory cues. Regardless, in both studies participants reported high confidence false percepts on noise-only trials in almost identical proportions (35% and 36% in the present and previous study, respectively). In the layer-specific imaging study, we found that the content of false percepts was reflected in the middle layers of the early visual cortex, suggesting a feedforward signal driving the content of false percepts. Together with the findings of the present study, we speculate that on noise-only trials, enhanced alpha/beta power and stimulus-specific activity fluctuations in the early visual cortex work in concert to generate high confidence false percepts. In line with this, increased alpha/beta power preceding false percepts source localised to a parieto-occipital network, making it ideally situated to interact with orientation-specific signals in the early visual cortex, potentially enhancing them through gain modulation. Indeed, previous studies have linked interactions between these areas to the modulation of sensory activity by priors (Rahnev et al., 2011). The nature of this potential interaction between alpha/beta oscillations and stimulus-specific activity fluctuations requires much further investigation.

Beyond a sensory signal reflecting the content of the reported false percepts, we also found a signal reflecting the confidence in stimulus presence, which emerged 250-300ms post-stimulus. Cross-decoding between grating-present and noise-only trials reveals that this neural signal was shared between false and veridical percepts. One might argue that this signal reflects motor preparation, potentially due to a facilitated motor response on high confidence trials. However, this is unlikely, because participants were required to provide an orientation response before their confidence response. Because the response-mapping for the orientation response was randomised, a motor response could not be prepared, precluding the possibility for the presence of a motor preparation signal. Instead, the uncovered neural signal is more likely to track confidence in the presence of a stimulus. Previous studies using variations of perceptual discrimination and detection tasks have reported neural correlates of perceptual confidence signals originating from the dorsolateral and (ventral) medial prefrontal cortex (Bang et al., 2020; Bang & Fleming, 2018; Gherman & Philiastides, 2018; Lau & Passingham, 2006; Shekhar & Rahnev, 2018; Yeon et al., 2020). Note that confidence here reflected the degree to which one was certain a stimulus was present, and is therefore strongly linked to stimulus visibility, which previous studies have shown to be encoded in frontal regions (Mazor et al., 2022). A likely neuromodulatory system that contributes to this signal is the dopamine system, which extensively innervates frontal regions (Björklund & Dunnett, 2007). Indeed, a series of studies have shown that dopamine may play an important role in modulating perceptual confidence, with dopaminergic agonists increasing confidence in perceptual detection (Lou et al., 2011). In rats, causally manipulating dopamine modulates confidence in false alarms (Schmack et al., 2021). Further, in primates, subjective stimulus intensity is reflected by the dopamine signal (De Lafuente & Romo, 2011). Most relevantly, a dopaminergic perceptual confidence signal in the caudate emerges around 300ms post-stimulus during decision-making (Lak et al., 2017), aligning with the emergence of the confidence signal in our cluster-based analyses. However, it should be noted that cluster-based analyses preclude exact inferences about the latency of signals (Maris & Oostenveld, 2007; Sassenhagen & Draschkow, 2019).

A number of influential studies have reported that false percepts are driven by overly strong priors (Schmack et al., 2021; Powers et al., 2017; Kafadar et al., 2020; Haarsma et al., 2020). While the false percepts in the present study were not induced by the perceptual expectations that we tried to elicit using predictive auditory cues, we do not think this should be taken as evidence against the predictive coding account of hallucinations. Rather, it highlights a possible additional mechanism through which prior expectations might induce false percepts. How might this work? In theory, perceptual priors could act on either level of stimulus content, or the level of stimulus presence. Indeed, both types of priors could lead to an increased tendency to report false alarms. For example, a cue signalling the onset of a sensory stimulus could increase false alarms by injecting sensory-like signals in the visual system. This could subsequently be mistaken for an external stimulus, as it passes a reality threshold, leading to the experience of a false percept. Indeed, recent work has suggested that imagination might mix with sensory signals, which under certain circumstances can lead to these signals to be mistaken for externally caused signals (Dijkstra et al., 2023). Alternatively, a prior that signals the likely presence of a stimulus, regardless of its content, could drive false percepts by increasing the likelihood of reporting a stimulus being present (Powers et al., 2017; Kafadar et al., 2020; Benrimoh et al., 2023). A study conducted in mice suggests that the dopamine system could be a candidate neurotransmitter system for acting as a prior on stimulus presence. Here, high confidence false percepts were related to, and could be induced by, increasing pre-stimulus dopamine levels in the caudate (Schmack et al., 2021). Indeed, previous work has linked dopamine in the caudate to perceptual confidence, thus increasing dopamine in this region pre-stimulus could de facto resemble a prior on stimulus presence (Lak et al., 2017). Alternatively, dopamine could mediate content-specific priors downstream. Regardless, it is clear that false percepts could arise in various ways. Ultimately, future studies are needed to study whether hallucination proneness is associated with an increased reliance on priors or presence, or priors on content, or both.

What could the role of increased alpha/beta power be in driving high confidence false percepts? As discussed above, in stimulus detection paradigms, it is well known that a decrease in alpha and beta power is related to increased hit rates (Achim et al., 2013; Benwell et al., 2017; Boncompte et al., 2016; Brüers & VanRullen, 2018; Chaumon & Busch, 2014; Ergenoglu et al., 2004; Mathewson et al., 2014; Samaha et al., 2017, 2020; Van Dijk et al., 2008). However, some studies have found the opposite for false percepts, where it is an increase in beta power, rather than a decrease, that drives false percepts (Keil et al., 2014; Poorganji et al., 2023). This suggests that the mechanism underlying perceptual thresholds for external stimuli (increased excitability) might be different from those for internally driven false percepts. One possible mechanism is suggested by previous studies which have linked high alpha/beta power to top-down endogenous signals. A number of studies have substantiated this, linking high alpha/beta power – often arising from the parietal cortex – to attention, imagination, illusory perception, and prediction (Arnal & Giraud, 2012; Auksztulewicz et al., 2017; Bartsch et al., 2015; Bastos et al., 2020; Buschman & Miller, 2007, 2009; De Lange et al., 2013; Engel & Fries, 2010; Fujioka et al., 2012; Hipp et al., 2011; Iversen et al., 2009; Okazaki et al., 2008; Pesaran et al., 2008; Raposo et al., 2023; Sherman et al., 2016; Villena-González et al., 2018). Moreover, previous fMRI studies have reported lower BOLD activity prior to false percepts (Hesselmann et al., 2010; Pajani et al., 2015). Because alpha/beta power and BOLD activity are inversely correlated (Scheeringa et al., 2011, 2016), speculatively, increased alpha/beta power might have played a role in these fMRI studies as well. However, this does not answer the question of what mechanism underlies fluctuations in beta power in the present study. Speculatively, they might reflect beliefs about stimulus presence. That is, while the specifically cued orientations (e.g., a low tone predicts a left-tilted grating) did not influence which orientation participants reported on high confidence false percepts trials, there was likely still a strong expectation of *a grating* (of either orientation) being present on each trial, as the participants were not explicitly told that there would be noise-only trials. This was confirmed by participants grossly underestimating the proportion of noise-only trials upon debriefing. As previous studies have linked an increase in beta power to stimulus predictions (Auksztulewicz et al., 2017; Bastos et al., 2020; Fujioka et al., 2012; Turner et al., 2023), we speculate here that beta power may reflect endogenous trial-by-trial fluctuations in stimulus expectation, which may have contributed to the high confidence false percepts in this study. Additionally, top-down mechanisms like imagination or choice history could have added to these effects (Dijkstra & Fleming, 2023; Urai et al., 2019).

In conclusion, the present study revealed that the neural signals underlying false and veridical perception are largely shared, both at the level of content-specific sensory signals and confidence in stimulus presence. Unique to false percepts, increased high alpha/low beta power arising from a parietal occipital network specifically preceded high confidence false percepts, but not veridical perception. Thereby, the current study sheds light on how false percepts arise from basic sensory mechanisms causing internal sensory signals to be mistaken for reality.

## Methods

### Ethics statement

This study was approved by the University College London Research Ethics Committee (R13061/RE002) and conducted according to the principles of the Declaration of Helsinki. All participants gave written informed consent prior to participation and received monetary compensation (£7.50 an hour for behavioural training, £10 an hour for MEG).

### Participants

Twenty-five healthy human volunteers with normal or corrected-to-normal vision participated in the MEG experiment. Two participants were excluded due to missing trigger data. The final sample consisted of 23 participants (22 female; age 25 ± 4 years; mean ± SD).

### Stimuli

Grayscale luminance-defined sinusoidal grating stimuli were generated using MATLAB (MathWorks, Natick, Massachusetts, United States of America, RRID:SCR_001622) and the Psychophysics Toolbox (Brainard, 1997). During the behavioural session, the stimuli were presented on a PC (1024 × 768 screen resolution, 60-Hz refresh rate). During the MEG recording session, stimuli were projected onto a screen in front of the participant (1920 × 1200 screen resolution, 60-Hz refresh rate). On grating-present trials (50%), auditory cues were followed by a grating after a 750ms delay (0.5-cpd spatial frequency, 33ms duration), displayed in an annulus (outer diameter: 10° of visual angle, inner diameter: 1°, contrast decreasing linearly to 0 over 0.7° at the inner and outer edges), surrounding a fixation bull’s eye (0.7° diameter). These stimuli were combined with one of 4 noise patches, which resulted in a 4% contrast grating embedded in 20% contrast noise during the MEG session. On noise-only trials, one of the 4 noise patches was presented on its own. Noise patches were created by smoothing pixel-by-pixel Gaussian noise with a Gaussian smoothing filter, ensuring that the spatial frequency of the noise matched the gratings (Wyart et al., 2012). This was done to ensure that the noise patches and gratings had similar low-level properties, increasing the likelihood of false percepts (Pajani et al., 2015). To avoid including noise patches which contained grating-like orientation signals by chance, 1000 noise patches were processed through a bank of Gabor filters with varying preferred orientations. Only noise patches with low (2%) signal energy for all orientations were selected to be included in the present experiment. The resulting four noise patches were used for all trials throughout the experiment, in a counterbalanced manner, ensuring that reported false percepts could only be triggered by internal mechanisms (Haarsma et al., 2023; Pajani et al., 2015). During the practice session on the first day, the contrast of the gratings was initially high (80%), gradually decreasing to 4% towards the end of the practice. The central fixation bull’s-eye was present throughout the trial, as well as during the intertrial interval (ITI; randomly varied between 1000 and 1200ms).

### Experimental procedure

Participant were required to perform a visual perceptual discrimination task. Trials consisted of an auditory expectation cue, followed by a grating stimulus embedded in noise on 50% of trials (750ms stimulus onset asynchrony (SOA) between cue and grating). The auditory cue (high or a low tone) predicted the orientation of the grating stimulus (45° or 135°) on grating-present trials. On these grating-present trials, a grating with the orientation predicted by the auditory cue was presented embedded in noise, while on noise-only trials (50%) only a noise patch was presented. The stimulus was presented for 33ms. After the stimulus disappeared, the orientation response prompt appeared, consisting of a left and right pointing arrow on either side of the fixation dot (location was counterbalanced). Participants were required to select the arrow corresponding to their answer (left arrow for anti-clockwise, or 135°, right arrow for clockwise, or 45°; 1s response window) through a button press with their right hand, using either a button box in the MEG, or a keyboard during behavioural training. Subsequently the letters “CONF?” appeared on the screen probing participants to indicate their confidence that they had seen a grating (1 = I did not see a grating, 2 = I may have seen a grating, 3 = I probably saw a grating, 4 = I am sure I saw a grating), using 1 of 4 buttons with their left hand (1.25s response window). It is worth highlighting here that confidence here reflected participants’ belief that a grating was present, not their confidence in their orientation report. It therefore reflects a perceptual awareness scale response (Sandberg & Overgaard, 2015). On the first day of testing, participants took part in a behavioural practice session. The practice consisted of an instruction phase with 7 blocks of 16 trials where the task was made progressively more difficult, whilst verbal and written instructions were provided. During these practice runs, the auditory cues predicted the orientation of the grating stimulus with 100% validity (45° or 135°; no noise-only trials). After the completion of the instructions, the participants completed 4 runs of 128 trials each, separated into 2 blocks of 64 trials each. In the first 2 runs the expectation cues were 100% valid, to ensure participants learnt the association, whilst in the final 2 runs the cues were 75% valid (i.e., the grating had an unexpected orientation on 25% of trials), to test whether participants might have adopted a response bias. Grating contrast decreased over the 4 runs, specifically the contrast levels were 7.5, 6, 5, and 4%, while the contrast of the noise patches remained constant at 20%. No noise-only trials were presented on day 1. On the second day, participants performed the same task during the MEG recording. During this session, 8 runs were completed, each consisting of 64 trials. This time the grating contrast was fixed at 4% on grating-present trials, and on 50% of the trials the gratings were omitted and only noise patches were presented, resulting in noise-only trials (Fig. 1a). On grating-present trials the cues always predicted the orientation of the grating with 100% validity (Fig. 1b). On noise-only trials the cue was by definition invalid, since no grating was presented. After each run a localiser run followed where gratings oriented either 45° or 135° were presented while participants performed a distracting fixation dimming task. The purpose of this localiser run was to uncover an orientation-specific MEG signal, which will not feature in the present paper as we were not able to uncover a significant orientation-specific signal evoked by the noisy stimuli presented in the current experiment. This analysis did reveal orientation-specific activity during the response window, which will be reported in detail elsewhere. Each run lasted ∼8 minutes, totalling ∼64 minutes.

### Pre-processing of MEG data

MEG was recorded continuously at 600 samples/second, using a whole-head 273 channel axial gradiometer system (CTF Omega, VSM MedTech), while participants sat upright. A photodiode was used to measure the onset of the visual stimuli through the presentation of a small white square in the bottom-right corner of the screen on both localiser trials as well as main experiment trials. This was done to ensure that the trials were aligned exactly to stimulus presentation. Note that the white square was not visible to participants, as it was covered by the electrode. The first experimental run (out of 8) for each subject was removed, leaving 7 runs for analyses. Trials were segmented 3000ms pre-stimulus and 3000ms post-stimulus. Movement and eye-blink artefacts were manually selected and inspected before being rejected from the data. Independent component analyses were applied to the complete dataset to identify components that reflect eye-blinks as well as cardiac related signals, which were manually inspected and removed from the data for each subject.

### ERF analyses

ERF analyses were conducted for exploratory reasons. We tested for differences in ERF amplitude for high and low confidence trials. Cluster-based analyses where then conducted on the sensor-level using Monte-Carlo permutation tests (N=10000), at a significance threshold of *p*<.05 for the initial threshold for determining a significant difference in ERF as well as for determining significant clusters. ERFs were computed separately for high and low confidence trials, for both grating-present and noise-only trials.

### Frequency analyses

Frequency power was estimated across all MEG channels from 2000ms pre-stimulus to 2000ms post-stimulus in steps of 50ms, for the frequencies 2Hz to 30Hz with steps of 1Hz, using Morlet wavelets (width=7). To test for pre-stimulus changes in alpha and beta power, we used cluster-based-permutation tests with 10000 iterations, at a significance threshold of *p* < 0.05, while averaging over the alpha band (8-12Hz) or the beta band (12-20Hz). For pre-stimulus effects we tested the time window from the onset of the cue to the onset of the stimulus, i.e. −750ms to 0ms. For post-stimulus effects the time window of interest ranged from 0ms to 1000ms. We conducted additional exploratory analyses that included the full pre-stimulus time-window −2000ms and full post-stimulus time-window. Note that these time windows include the onset of the auditory stimulus (−750ms) and the orientation response cues (1000ms). We furthermore repeated the analyses for each frequency between 8 and 20 Hz with steps of 1Hz to estimate which specific frequencies drove the effects.

### Decoding analyses

We decoded both stimulus content (grating orientation) as well as participants’ confidence in having seen a stimulus from the MEG signal using a two-class linear discriminant analysis (LDA) (Mostert et al. 2015). For content decoding, we were interested in testing whether subjectively perceived orientations were accompanied by sensory-like orientation signals, on both false percept and veridical trials. To obtain a measure of a sensory orientation signal, we trained a decoder on MEG data from separate localiser blocks to distinguish the orientation of clearly presented (but task-irrelevant) gratings. Localiser blocks were presented at the end of each run (of which there were 8), which each contained 80 trials of clearly presented gratings. Once trained, we applied this decoder to the main experiment, to test for orientation-specific neural activity on false percept and veridical trials. To train the decoder, the 273 MEG channels were used as features. To confirm the ability of the decoder to retrieve orientation signals, we also trained and tested it within the localiser data using a leave-one out procedure, where all localiser blocks except one served as the training data to decode the remaining block. This procedure was repeated for all blocks. Time points were averaged across a moving time window containing 17 samples ((17/600) = 28.3ms), with steps of 3 samples ((3/600) = 5ms). The covariance matrix was taken into account in the decoder as recommended in previous studies to address correlations between neighbouring MEG sensors (Brouwer & Heeger, 2009; Mostert et al., 2015). For details on the implementation of the decoder, see Mostert et al. 2015.

For the purpose of decoding confidence in having seen a stimulus, we used a cross-validated decoding procedure similar to the approach used to decode orientations within the localiser blocks. Specifically, using a leave-one-block-out procedure, we decoded confidence in having seen a grating during the main experiment, by training the LDA on binarized confidence response, such that the confidence responses reflected higher or lower than average confidence. All other parameters of the LDA were identical to the orientation decoding analysis.

To test for sensory-like orientation-specific activity during the main experiment we trained on the group-level peak of the localiser decoding and averaged over a 10ms time window (100ms-110ms), and generalised to the main experiment. Subsequently, the evidence traces for trials where a left-tilted grating was reported was subtracted from right-tilted traces, resulting in relative evidence in favour of the reported stimulus. Subsequently, low confidence trials were subtracted from high confidence trials to show activity specific for orientations that the participant indicated to have seen. Then, cluster-based permutation tests were conducted on the time courses of these signals. In the first step of the permutation test, clusters were defined by adjacent points that crossed a significance threshold of *p* <.05. The number of permutations for the temporal generalisation matrix was limited to 1,000 due to the large sample space. In contrast, when performing cluster-based analyses on a single time course, as when we generalised from the localiser to the main experiment, or the diagonal of the temporal generalisation matrix for the confidence decoding 10,000 permutations were used. A cluster in the true data was considered significant if *p*<.05 based on the null distribution generated by the permutations. The time window for the confidence decoding was initially −1000ms to 2000ms relative to stimulus onset to include the time window where the orientation responses were given (1000-2000ms). For the analyses pertaining to finding content specific signals, we focussed on pre- and post-stimulus time windows separately, and excluded the time window where the orientation was reported (i.e., 1000ms onwards). We performed a number of follow-up analyses for the confidence decoding. These follow-up analyses involved cross-decoding between noise-only and grating-present trials, to explore whether they shared a neural code for perceptual confidence. Furthermore, we performed control analyses where we balanced the number of trials in the high and low perceptual confidence conditions by subsampling to the condition with the fewest trials, to ensure that the effects were not driven by an overrepresentation of a single condition.

### Source localisation

To visualise the source of the confidence decoding we performed source localisation analyses. We did not collect individual anatomical MRI scans for our subjects, instead, a template MRI and default head and source models as present in the Fieldtrip toolbox (www.fieldtriptoolbox.org) were used (see below) (Oostenveld et al., 2011). Previous studies have demonstrated that little anatomical specificity is lost using a group-based template approach (Holliday et al., 2003).

The spatial pattern that underlies classification in a linear discrimination analysis is driven by the difference in magnetic fields between the two conditions on which the decoding is based. Therefore, one can visualise the source of a decoder by estimating the sources of the two different conditions, and compute the difference (Haufe et al., 2014). For our purposes we computed the absolute difference within the 1000ms to 2000ms time window (where decoding was strongest), divided by the source map of the low confidence condition, thereby estimating percentage signal change. Taking the absolute difference will visualise which source signals are involved in perceptual confidence, without making assumptions about the sign of the signal.

We also performed source localisation on the frequency analyses. Here, there was a direct translation from the univariate effects to the underlying source map, as they seek to refute the same null-hypothesis. We therefore computed cluster-based staisics on the −750ms to 0ms ime window for the difference in 13Hz (+- 1.5Hz) power between high and low confidence false percept condiions in source space. The same parameters were used as on the sensor level (10000 permutations, cluster-defining threshold: *p*=.05, cluster-level threshold: *p*=.05).

For both the frequency and decoding source localisation, we used the default forward and source models from the Fieldtrip toolbox which were then warped to participants’ specific fiducials based on the MEG sensors. The spatial filter was computed for the time windows of interest in the averaged data, which was subsequently applied separately to the two conditions of interest (high and low confidence trials). Source localisation was then computed for the two conditions of interests. For the confidence decoding a percentage absolute signal change was computed in source space, whereas for the frequency analyses t-maps were visualised for significant clusters.

### Behavioural analyses

We tested participants’ accuracy scores, as well as the relationship between accuracy and confidence in stimulus presence. We further tested whether participants were significantly biased by the auditory cues on noise-only trials, and performed post-hoc tests to see whether these effects were driven by cue awareness. All tests were conducted within-subject and two-sided. Statistics were conducted in JASP (JASP team, 2023).

## Acknowledgements

This work was supported by a Wellcome/Royal Society Sir Henry Dale Fellowship [218535/Z/19/Z] and a European Research Council (ERC) Starting Grant [948548] to P.K. The Wellcome Centre for Human Neuroimaging is supported by core funding from the Wellcome Trust [203147/Z/16/Z].

---

**Supplementary Fig. 1.**
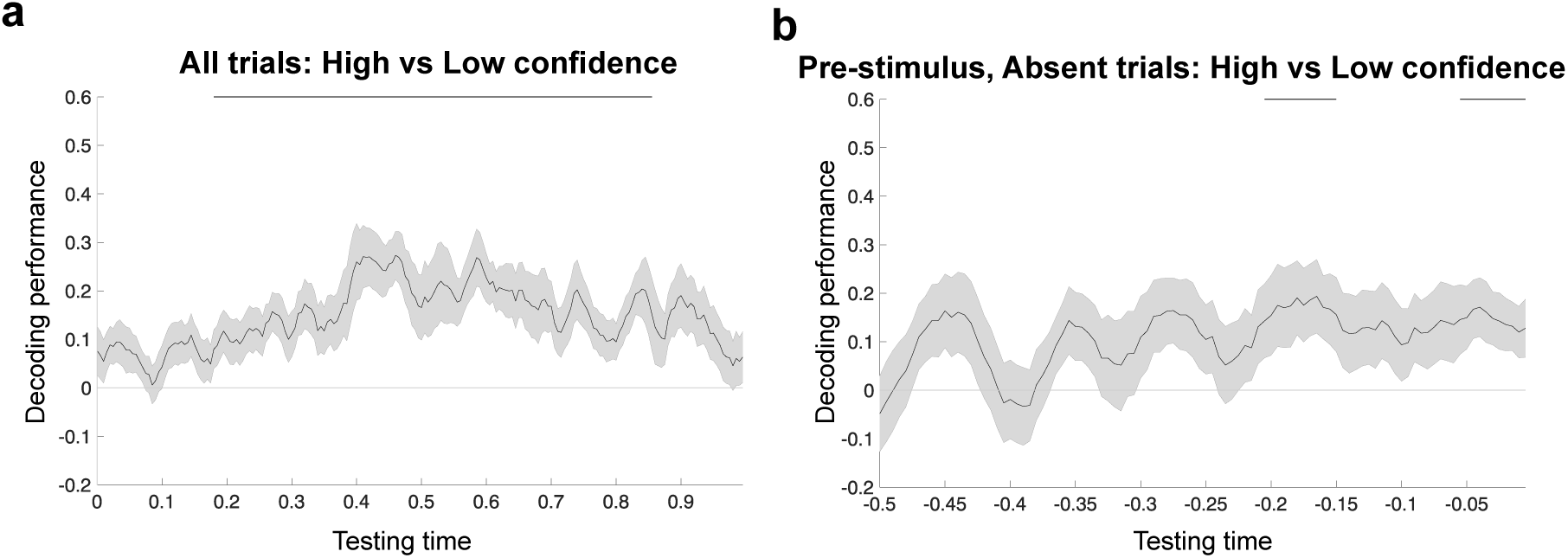
Additional figures for orientation and stimulus-specific decoding of perceived gratings. **a,** Subtracting low from high confidence false percepts, a clear signal is present reflecting the orientation of perceived gratings on all trials. **b,** On false percept trials there was additional orientation and stimulus-specific activity reflecting false percepts in the pre-stimulus time window.

**Supplementary Fig. 2.**
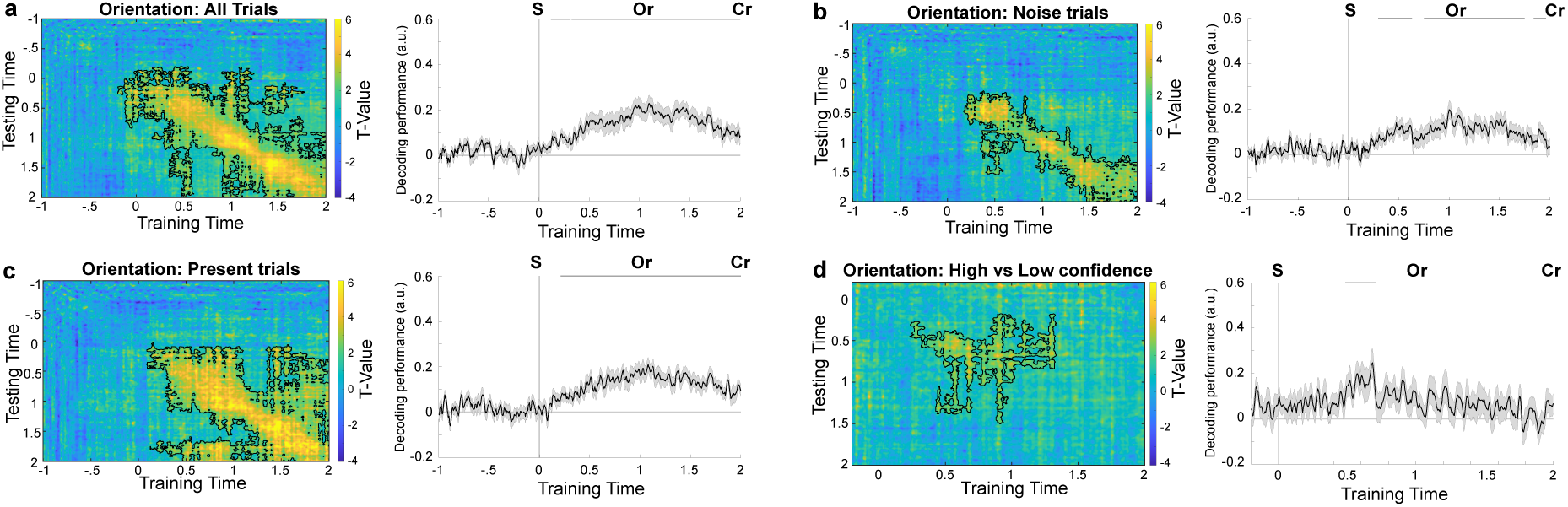
Within decoding of reported orientation. **a,** As a comparison to the within confidence decoding we used the same approach to decode the reported orientations during the main experiment. Taking all trials together, we find a signal reflecting the reported percept emerged from 120ms to 315ms post-stimulus (*p*=.026, Cohen’s d=1.02), and from 325ms until the end of the decoding time window of 2000ms (*p*<.001, Cohen’s d=1.13). **b,** This signal was present on both noise-only (295-630ms, 750-1750ms & 1840-1960ms *p*=.0018, *p*<.001 & *p*=.038, Cohen’s d= 1.03, .89 & .65 respectively). **c,** and grating-present trials (210ms-2000ms, *p*<.001, Cohen’s d= 1.43). **d,** This signal was modulated by participants confidence in stimulus presence, such that decoding was stronger for high confidence trials at 485-705ms, which overlaps with the timewindow for sensory-like signals that underlie false percepts (*p*=.0019, Cohen’s d = .77).

## References

Achim, A., Bouchard, J., & Braun, C. M. J. (2013). EEG amplitude spectra before near threshold visual presentations differentially predict detection/omission and short– long reaction time outcomes. International Journal of Psychophysiology, 89(1), 88–98. 10.1016/j.ijpsycho.2013.05.016

Aitken, F., Menelaou, G., Warrington, O., Koolschijn, R. S., Corbin, N., Callaghan, M. F., & Kok, P. (2020). Prior expectations evoke stimulus-specific activity in the deep layers of the primary visual cortex. PLOS Biology, 18(12), e3001023. 10.1371/journal.pbio.3001023

Albers, A. M., Kok, P., Toni, I., Dijkerman, H. C., & de Lange, F. P. (2013). Shared Representations for Working Memory and Mental Imagery in Early Visual Cortex. Current Biology, 23(15), 1427–1431. 10.1016/j.cub.2013.05.065

Arnal, L. H., & Giraud, A.-L. (2012). Cortical oscillations and sensory predictions. Trends in Cognitive Sciences, 16(7), 390–398. 10.1016/j.tics.2012.05.003

Auksztulewicz, R., Friston, K. J., & Nobre, A. C. (2017). Task relevance modulates the behavioural and neural effects of sensory predictions. PLOS Biology, 15(12), e2003143. 10.1371/journal.pbio.2003143

Bang, D., Ershadmanesh, S., Nili, H., & Fleming, S. M. (2020). Private–public mappings in human prefrontal cortex. eLife, 9, e56477. 10.7554/eLife.56477

Bang, D., & Fleming, S. M. (2018). Distinct encoding of decision confidence in human medial prefrontal cortex. Proceedings of the National Academy of Sciences, 115(23), 6082– 6087. 10.1073/pnas.1800795115

Bartsch, F., Hamuni, G., Miskovic, V., Lang, P. J., & Keil, A. (2015). Oscillatory brain activity in the alpha range is modulated by the content of word-prompted mental imagery: Alpha oscillations during mental imagery. Psychophysiology, 52(6), 727–735. 10.1111/psyp.12405

Bastos, A. M., Lundqvist, M., Waite, A. S., Kopell, N., & Miller, E. K. (2020). Layer and rhythm specificity for predictive routing. Proceedings of the National Academy of Sciences, 117(49), 31459–31469. 10.1073/pnas.2014868117

Bastos, A. M., Usrey, W. M., Adams, R. A., Mangun, G. R., Fries, P., & Friston, K. J. (2012). Canonical Microcircuits for Predictive Coding. Neuron, 76(4), 695–711. 10.1016/j.neuron.2012.10.038

Benwell, C. S. Y., Tagliabue, C. F., Veniero, D., Cecere, R., Savazzi, S., & Thut, G. (2017). Prestimulus EEG Power Predicts Conscious Awareness But Not Objective Visual Performance. Eneuro, 4(6), ENEURO.0182-17.2017. 10.1523/ENEURO.0182-17.2017

Björklund, A., & Dunnett, S. B. (2007). Dopamine neuron systems in the brain: An update. Trends in Neurosciences, 30(5), 194–202. 10.1016/j.tins.2007.03.006

Boncompte, G., Villena-González, M., Cosmelli, D., & López, V. (2016). Spontaneous Alpha Power Lateralization Predicts Detection Performance in an Un-Cued Signal Detection Task. PLOS ONE, 11(8), e0160347. 10.1371/journal.pone.0160347

Brouwer, G. J., & Heeger, D. J. (2009). Decoding and Reconstructing Color from Responses in Human Visual Cortex. The Journal of Neuroscience, 29(44), 13992–14003. 10.1523/JNEUROSCI.3577-09.2009

Brüers, S., & VanRullen, R. (2018). Alpha Power Modulates Perception Independently of Endogenous Factors. Frontiers in Neuroscience, 12, 279. 10.3389/fnins.2018.00279

Buschman, T. J., & Miller, E. K. (2007). Top-Down Versus Bottom-Up Control of Attention in the Prefrontal and Posterior Parietal Cortices. Science, 315(5820), 1860–1862. 10.1126/science.1138071

Buschman, T. J., & Miller, E. K. (2009). Serial, Covert Shifts of Attention during Visual Search Are Reflected by the Frontal Eye Fields and Correlated with Population Oscillations. Neuron, 63(3), 386–396. 10.1016/j.neuron.2009.06.020

Chaumon, M., & Busch, N. A. (2014). Prestimulus Neural Oscillations Inhibit Visual Perception via Modulation of Response Gain. Journal of Cognitive Neuroscience, 26(11), 2514–2529. 10.1162/jocn_a_00653

Collerton, D., Barnes, J., Diederich, N. J., Dudley, R., Ffytche, D., Friston, K., Goetz, C. G., Goldman, J. G., Jardri, R., Kulisevsky, J., Lewis, S. J. G., Nara, S., O’Callaghan, C., Onofrj, M., Pagonabarraga, J., Parr, T., Shine, J. M., Stebbins, G., Taylor, J.-P., … Weil, R. S. (2023). Understanding visual hallucinations: A new synthesis. Neuroscience & Biobehavioral Reviews, 150, 105208. 10.1016/j.neubiorev.2023.105208

De Lafuente, V., & Romo, R. (2011). Dopamine neurons code subjective sensory experience and uncertainty of perceptual decisions. Proceedings of the National Academy of Sciences, 108(49), 19767–19771. 10.1073/pnas.1117636108

De Lange, F. P., Rahnev, D. A., Donner, T. H., & Lau, H. (2013). Prestimulus Oscillatory Activity over Motor Cortex Reflects Perceptual Expectations. The Journal of Neuroscience, 33(4), 1400–1410. 10.1523/JNEUROSCI.1094-12.2013

Dijkstra, N., Bosch, S. E., & Van Gerven, M. A. J. (2017). Vividness of Visual Imagery Depends on the Neural Overlap with Perception in Visual Areas. The Journal of Neuroscience, 37(5), 1367–1373. 10.1523/JNEUROSCI.3022-16.2016

Dijkstra, N., & Fleming, S. M. (2023). Subjective signal strength distinguishes reality from imagination. Nature Communications, 14(1), 1627. 10.1038/s41467-023-37322-1

Dijkstra, N., Mostert, P., Lange, F. P. D., Bosch, S., & Van Gerven, M. A. (2018). Differential temporal dynamics during visual imagery and perception. eLife, 7, e33904. 10.7554/eLife.33904

Engel, A. K., & Fries, P. (2010). Beta-band oscillations—Signalling the status quo? Current Opinion in Neurobiology, 20(2), 156–165. 10.1016/j.conb.2010.02.015

Engel, A. K., Fries, P., & Singer, W. (2001). Dynamic predictions: Oscillations and synchrony in top–down processing. Nature Reviews Neuroscience, 2(10), 704–716. 10.1038/35094565

Ergenoglu, T., Demiralp, T., Bayraktaroglu, Z., Ergen, M., Beydagi, H., & Uresin, Y. (2004). Alpha rhythm of the EEG modulates visual detection performance in humans. Cognitive Brain Research, 20(3), 376–383. 10.1016/j.cogbrainres.2004.03.009

Fujioka, T., Trainor, L. J., Large, E. W., & Ross, B. (2012). Internalized Timing of Isochronous Sounds Is Represented in Neuromagnetic Beta Oscillations. The Journal of Neuroscience, 32(5), 1791–1802. 10.1523/JNEUROSCI.4107-11.2012

Gherman, S., & Philiastides, M. G. (2018). Human VMPFC encodes early signatures of confidence in perceptual decisions. eLife, 7, e38293. 10.7554/eLife.38293

Haarsma, J., Deveci, N., Corbin, N., Callaghan, M. F., & Kok, P. (2022). *Perceptual expectations and false percepts generate stimulus-specific activity in distinct layers of the early visual cortex* [Preprint]. Neuroscience. 10.1101/2022.04.13.488155

Haarsma, J., Deveci, N., Corbin, N., Callaghan, M. F., & Kok, P. (2023). Expectation cues and false percepts generate stimulus-specific activity in distinct layers of the early visual cortex Laminar profile of visual false percepts. The Journal of Neuroscience, JN-RM-0998–23. 10.1523/JNEUROSCI.0998-23.2023

Harrison, S. A., & Tong, F. (2009). Decoding reveals the contents of visual working memory in early visual areas. Nature, 458(7238), 632–635. 10.1038/nature07832

Haufe, S., Meinecke, F., Görgen, K., Dähne, S., Haynes, J.-D., Blankertz, B., & Bießmann, F. (2014). On the interpretation of weight vectors of linear models in multivariate neuroimaging. NeuroImage, 87, 96–110. 10.1016/j.neuroimage.2013.10.067

Hesselmann, G., Sadaghiani, S., Friston, K. J., & Kleinschmidt, A. (2010). Predictive Coding or Evidence Accumulation? False Inference and Neuronal Fluctuations. PLoS ONE, 5(3), e9926. 10.1371/journal.pone.0009926

Hipp, J. F., Engel, A. K., & Siegel, M. (2011). Oscillatory Synchronization in Large-Scale Cortical Networks Predicts Perception. Neuron, 69(2), 387–396. 10.1016/j.neuron.2010.12.027

Holliday, I. E., Barnes, G. R., Hillebrand, A., & Singh, K. D. (2003). Accuracy and applications of group MEG studies using cortical source locations estimated from participants’ scalp surfaces. Human Brain Mapping, 20(3), 142–147. 10.1002/hbm.10133

Iemi, L., & Busch, N. A. (2018). Moment-to-Moment Fluctuations in Neuronal Excitability Bias Subjective Perception Rather than Strategic Decision-Making. Eneuro, 5(3), ENEURO.0430-17.2018. 10.1523/ENEURO.0430-17.2018

Iemi, L., Chaumon, M., Crouzet, S. M., & Busch, N. A. (2017). Spontaneous Neural Oscillations Bias Perception by Modulating Baseline Excitability. The Journal of Neuroscience, 37(4), 807–819. 10.1523/JNEUROSCI.1432-16.2016

Iversen, J. R., Repp, B. H., & Patel, A. D. (2009). Top-Down Control of Rhythm Perception Modulates Early Auditory Responses. Annals of the New York Academy of Sciences, 1169(1), 58–73. 10.1111/j.1749-6632.2009.04579.x

Keil, J., Müller, N., Hartmann, T., & Weisz, N. (2014). Prestimulus Beta Power and Phase Synchrony Influence the Sound-Induced Flash Illusion. Cerebral Cortex, 24(5), 1278– 1288. 10.1093/cercor/bhs409

Kok, P., Failing, M. F., & de Lange, F. P. (2014). Prior Expectations Evoke Stimulus Templates in the Primary Visual Cortex. Journal of Cognitive Neuroscience, 26(7), 1546–1554. 10.1162/jocn_a_00562

Kok, P., Mostert, P., & De Lange, F. P. (2017). Prior expectations induce prestimulus sensory templates. Proceedings of the National Academy of Sciences, 114(39), 10473–10478. 10.1073/pnas.1705652114

Lak, A., Nomoto, K., Keramati, M., Sakagami, M., & Kepecs, A. (2017). Midbrain Dopamine Neurons Signal Belief in Choice Accuracy during a Perceptual Decision. Current Biology, 27(6), 821–832. 10.1016/j.cub.2017.02.026

Lange, J., Oostenveld, R., & Fries, P. (2013). Reduced Occipital Alpha Power Indexes Enhanced Excitability Rather than Improved Visual Perception. The Journal of Neuroscience, 33(7), 3212–3220. 10.1523/JNEUROSCI.3755-12.2013

Lau, H. C., & Passingham, R. E. (2006). Relative blindsight in normal observers and the neural correlate of visual consciousness. Proceedings of the National Academy of Sciences, 103(49), 18763–18768. 10.1073/pnas.0607716103

Lawrence, S. J. D., Van Mourik, T., Kok, P., Koopmans, P. J., Norris, D. G., & De Lange, F. P. (2018). Laminar Organization of Working Memory Signals in Human Visual Cortex. Current Biology, 28(21), 3435–3440.e4. 10.1016/j.cub.2018.08.043

Lou, H. C., Skewes, J. C., Thomsen, K. R., Overgaard, M., Lau, H. C., Mouridsen, K., & Roepstorff, A. (2011). Dopaminergic stimulation enhances confidence and accuracy in seeing rapidly presented words. Journal of Vision, 11(2), 15–15. 10.1167/11.2.15

Maris, E., & Oostenveld, R. (2007). Nonparametric statistical testing of EEG- and MEG-data. Journal of Neuroscience Methods, 164(1), 177–190. 10.1016/j.jneumeth.2007.03.024

Mathewson, K. E., Beck, D. M., Ro, T., Maclin, E. L., Low, K. A., Fabiani, M., & Gratton, G. (2014). Dynamics of Alpha Control: Preparatory Suppression of Posterior Alpha Oscillations by Frontal Modulators Revealed with Combined EEG and Event-related Optical Signal. Journal of Cognitive Neuroscience, 26(10), 2400–2415. 10.1162/jocn_a_00637

Mazor, M., Dijkstra, N., & Fleming, S. M. (2022). Dissociating the Neural Correlates of Subjective Visibility from Those of Decision Confidence. The Journal of Neuroscience, 42(12), 2562–2569. 10.1523/JNEUROSCI.1220-21.2022

Mostert, P., Kok, P., & De Lange, F. P. (2015). Dissociating sensory from decision processes in human perceptual decision making. Scientific Reports, 5(1), 18253. 10.1038/srep18253

Okazaki, M., Kaneko, Y., Yumoto, M., & Arima, K. (2008). Perceptual change in response to a bistable picture increases neuromagnetic beta-band activities. Neuroscience Research, 61(3), 319–328. 10.1016/j.neures.2008.03.010

Oostenveld, R., Fries, P., Maris, E., & Schoffelen, J.-M. (2011). FieldTrip: Open Source Software for Advanced Analysis of MEG, EEG, and Invasive Electrophysiological Data. Computational Intelligence and Neuroscience, 2011, 1–9. 10.1155/2011/156869

Pajani, A., Kok, P., Kouider, S., & de Lange, F. P. (2015). Spontaneous Activity Patterns in Primary Visual Cortex Predispose to Visual Hallucinations. Journal of Neuroscience, 35(37), 12947–12953. 10.1523/JNEUROSCI.1520-15.2015

Peelen, M. V., & Kastner, S. (2011). A neural basis for real-world visual search in human occipitotemporal cortex. Proceedings of the National Academy of Sciences, 108(29), 12125–12130. 10.1073/pnas.1101042108

Pesaran, B., Nelson, M. J., & Andersen, R. A. (2008). Free choice activates a decision circuit between frontal and parietal cortex. Nature, 453(7193), 406–409. 10.1038/nature06849

Poorganji, M., Zomorrodi, R., Zrenner, C., Bansal, A., Hawco, C., Hill, A. T., Hadas, I., Rajji, T. K., Chen, R., Zrenner, B., Voineskos, D., Blumberger, D. M., & Daskalakis, Z. J. (2023). Pre-Stimulus Power but Not Phase Predicts Prefrontal Cortical Excitability in TMS-EEG. Biosensors, 13(2), 220. 10.3390/bios13020220

Rahnev, D., Lau, H., & De Lange, F. P. (2011). Prior Expectation Modulates the Interaction between Sensory and Prefrontal Regions in the Human Brain. Journal of Neuroscience, 31(29), 10741–10748. 10.1523/JNEUROSCI.1478-11.2011

Raposo, I., Szczepanski, S. M., Haaland, K., Endestad, T., Solbakk, A.-K., Knight, R. T., & Helfrich, R. F. (2023). Periodic attention deficits after frontoparietal lesions provide causal evidence for rhythmic attentional sampling. Current Biology, S0960982223013143. 10.1016/j.cub.2023.09.065

Samaha, J., Iemi, L., Haegens, S., & Busch, N. A. (2020). Spontaneous Brain Oscillations and Perceptual Decision-Making. Trends in Cognitive Sciences, 24(8), 639–653. 10.1016/j.tics.2020.05.004

Samaha, J., Iemi, L., & Postle, B. R. (2017). Prestimulus alpha-band power biases visual discrimination confidence, but not accuracy. Consciousness and Cognition, 54, 47–55. 10.1016/j.concog.2017.02.005

Sánchez-Fuenzalida, N., Van Gaal, S., Fleming, S. M., Haaf, J. M., & Fahrenfort, J. J. (2023). Predictions and rewards affect decision-making but not subjective experience. Proceedings of the National Academy of Sciences, 120(44), e2220749120. 10.1073/pnas.2220749120

Sandberg, K., & Overgaard, M. (2015). Using the perceptual awareness scale (PAS). Behavioral Methods in Consciousness Research, 181–196.

Sassenhagen, J., & Draschkow, D. (2019). Cluster-based permutation tests of MEG/EEG data do not establish significance of effect latency or location. Psychophysiology, 56(6), e13335. 10.1111/psyp.13335

Scheeringa, R., Fries, P., Petersson, K.-M., Oostenveld, R., Grothe, I., Norris, D. G., Hagoort, P., & Bastiaansen, M. C. M. (2011). Neuronal Dynamics Underlying High- and Low-Frequency EEG Oscillations Contribute Independently to the Human BOLD Signal. Neuron, 69(3), 572–583. 10.1016/j.neuron.2010.11.044

Scheeringa, R., Koopmans, P. J., Van Mourik, T., Jensen, O., & Norris, D. G. (2016). The relationship between oscillatory EEG activity and the laminar-specific BOLD signal. Proceedings of the National Academy of Sciences, 113(24), 6761–6766. 10.1073/pnas.1522577113

Schmack, K., Bosc, M., Ott, T., Sturgill, J. F., & Kepecs, A. (2021). Striatal dopamine mediates hallucination-like perception in mice. Science, 372(6537), eabf4740. 10.1126/science.abf4740

Shekhar, M., & Rahnev, D. (2018). Distinguishing the Roles of Dorsolateral and Anterior PFC in Visual Metacognition. The Journal of Neuroscience, 38(22), 5078–5087. 10.1523/JNEUROSCI.3484-17.2018

Sherman, M. T., Kanai, R., Seth, A. K., & VanRullen, R. (2016). Rhythmic Influence of Top– Down Perceptual Priors in the Phase of Prestimulus Occipital Alpha Oscillations. Journal of Cognitive Neuroscience, 28(9), 1318–1330. 10.1162/jocn_a_00973

Stokes, M., Thompson, R., Cusack, R., & Duncan, J. (2009). Top-Down Activation of Shape-Specific Population Codes in Visual Cortex during Mental Imagery. The Journal of Neuroscience, 29(5), 1565–1572. 10.1523/JNEUROSCI.4657-08.2009

Stokes, M., Thompson, R., Nobre, A. C., & Duncan, J. (n.d.). Shape-specific preparatory activity mediates attention to targets in human visual cortex.

Turner, W., Blom, T., & Hogendoorn, H. (2023). Visual Information Is Predictively Encoded in Occipital Alpha/Low-Beta Oscillations. The Journal of Neuroscience, 43(30), 5537– 5545. 10.1523/JNEUROSCI.0135-23.2023

Urai, A. E., De Gee, J. W., Tsetsos, K., & Donner, T. H. (2019). Choice history biases subsequent evidence accumulation. eLife, 8, e46331. 10.7554/eLife.46331

Van Dijk, H., Schoffelen, J.-M., Oostenveld, R., & Jensen, O. (2008). Prestimulus Oscillatory Activity in the Alpha Band Predicts Visual Discrimination Ability. The Journal of Neuroscience, 28(8), 1816–1823. 10.1523/JNEUROSCI.1853-07.2008

Villena-González, M., Palacios-García, I., Rodríguez, E., & López, V. (2018). Beta Oscillations Distinguish Between Two Forms of Mental Imagery While Gamma and Theta Activity Reflects Auditory Attention. Frontiers in Human Neuroscience, 12, 389. 10.3389/fnhum.2018.00389

Wyart, V., Nobre, A. C., & Summerfield, C. (2012). Dissociable prior influences of signal probability and relevance on visual contrast sensitivity. Proceedings of the National Academy of Sciences, 109(9), 3593–3598. 10.1073/pnas.1120118109

Yeon, J., Shekhar, M., & Rahnev, D. (2020). Overlapping and unique neural circuits are activated during perceptual decision making and confidence. Scientific Reports, 10(1), 20761. 10.1038/s41598-020-77820-6

